# Unveiling the pathogenicity of *Cunninghamella echinulata* SMH-1 and its bioactive metabolites through metabolomics: Dual-action disruption of *Solenopsis invicta* mitochondrial bioenergetics and cuticular integrity

**DOI:** 10.1101/2025.04.10.648148

**Authors:** Mehboob Hussain, Xi Gao, Guoxing Wu, Deqiang Qin

## Abstract

The invasive red imported fire ant, *Solenopsis invicta*, poses significant ecological and economic challenges, necessitating sustainable alternatives to conventional pest control. This study evaluates the biocontrol potential of *Cunninghamella echinulata* isolate SMH-1, an entomopathogenic fungus, through integrated morphological, biochemical, and metabolomic analyses. Light and scanning electron microscopy elucidated infection dynamics, revealing conidial germination, cuticular penetration, and systemic hyphal proliferation, culminating in complete cadaver colonization. At 1×10⁸ conidia/mL, SMH-1 achieved 100% mortality in *S. invicta* workers within 7 days, with virulence maintained across thermal gradients (10–30°C). Acetone extracts of SMH- 1 induced dose-dependent lethality, reaching 100% mortality at 1,000 mg/L. Nineteen significantly differentiated metabolites were identified from phospholipids category through UPLC/MS analysis of isolate SMH-1. The derived compounds also showed strong potential against *S. invicta*, among which dihydrocoumarin caused 100% mortality over 12-days of exposure. The bioactivity of isolate SMH-1 (Conidia), (extracts) and (dihydrocoumarin) against *S. invicta* revealed significantly differentiated metabolites as (Sulfolithocholylglycine, PE(18_1(9Z)_0_0), CE(12:0), LysoPI(18:1(9Z)/0:0) and Hydroxydestruxin B), (Tropine, lignoceric acid, 9-deoxy-9-methylene-16,16-dimethyl-PGE2, Flecainide and Jubanine B), and (3-O-alpha-D-Glucopyranuronosyl-D-xylose, Rilmakalim, 2-Amino-3-methyl-1-butanol, 2,4-Dimethyl-1,3-oxazole-5-carboxylic acid and Caffeinol), respectively. Citrate cycle (TCA) was significantly impacted by differential expression of pyruvate, pyruvic acid, succinic acid and fumaric acid. These findings underscore SMH-1’s multifaceted biocontrol potential, combining direct pathogenicity, thermo-tolerance, and metabolite-mediated toxicity. By delineating host-pathogen interactions and metabolic disruptions, this work advances *C. echinulata* isolate SMH-1 as a sustainable candidate for integrated *S. invicta* management, offering an eco-friendly alternative to synthetic insecticides.

**Author Summary:** Invasive fire ants (*S. invicta*) threaten ecosystems and agriculture worldwide, and current chemical controls often harm the environment. In this study, we explored a natural alternative: a soil-dwelling fungus called *Cunninghamella echinulata* (strain SMH-1). We discovered that this fungus can kill fire ants with remarkable efficiency. When applied directly, its spores caused 100% ant mortality within a week, even under varying temperatures. We also identified key fungal compounds, including dihydrocoumarin, which are lethal to ants. By studying how the fungus invades and disrupts the ants’ biology-such as breaking down their protective outer layers and interfering with their energy production, we revealed why SMH-1 is so effective. Our findings highlight SMH-1’s potential as a safe, eco-friendly biocontrol agent. This fungus could reduce reliance on synthetic pesticides, offering farmers and communities a sustainable tool to combat fire ant invasions while protecting biodiversity. Our work bridges lab discoveries and real-world pest management, paving the way for greener solutions in agriculture and ecosystem conservation.

## 1. Introduction

Invasive alien species pose profound threats to agriculture, human health, native ecosystems, and global biodiversity^1–3^. Among these, the red imported fire ant (*Solenopsis invicta* Buren; Hymenoptera: Formicidae, RIFA) stands out as a highly aggressive and ecologically disruptive species, renowned for its rapid range expansion and colonization of diverse habitats, including agricultural lands, urban green spaces, and infrastructure^4–6^. Beyond ecological disruption, RIFA inflicts painful stings that endanger human health and necessitates costly public health interventions, while it’s nesting behavior damages electrical systems and irrigation networks^7^. Cumulative economic losses attributed to RIFA invasions in Australia and the United States alone exceed $10.95 billion USD (1930–2020), with 80% of these costs directly linked to its management^8^. Consequently, RIFA is now classified among the world’s 100 most damaging invasive species^5,9^, recognized as a global quarantine pest^10^, and recently designated by the European Union as a priority invasive alien species of concern^11^.

Current RIFA management relies heavily on synthetic insecticides, including pyrethroids, insect growth regulators (e.g., ivermectin), and contact neurotoxins such as chlorpyrifos^12,13^. However, prolonged reliance on chemical controls has led to environmental contamination, non-target organism mortality, and the emergence of resistance in RIFA populations^14,15^. This underscores the urgent need for sustainable alternatives within integrated pest management (IPM) frameworks. Entomopathogenic biocontrol agents-including bacteria, nematodes, and fungi-have gained prominence as ecologically compatible solutions^12^. Fungal pathogens, in particular, demonstrate high specificity and self-dispersing potential. Laboratory studies confirm the efficacy of *Beauveria bassiana*, *Metarhizium anisopliae*, and *M. flavoviride* against RIFA^16–20^, yet field applications remain constrained by environmental sensitivity and variable virulence. Thus, identifying novel fungal species with enhanced adaptability and pathogenicity is critical to advancing biocontrol strategies.

The genus *Cunninghamella* (Mucorales: Cunninghamellaceae), first described by Matruchot in 1903, comprises 14 recognized species characterized by unispored sporangia and pedicellate vesicles^21–23^. While commonly isolated from soil, stored grains, and decomposing organic matter^24,25^, its entomopathogenic potential remains largely unexplored. Notably, no prior studies have investigated *C. echinulata*’s virulence against RIFA, despite its phylogenetic proximity to established entomopathogens.

Entomopathogenic fungi typically infect hosts via cuticular penetration, mediated by hydrophobic interactions and enzymatic degradation of chitinous exoskeletons^26^. Concurrently, they produce secondary metabolites including terpenoids, cyclic depsipeptides, and polyketides-that suppress host immune responses and accelerate mortality^27,28^. These mycotoxins, often low-molecular-weight compounds, have demonstrated insecticidal activity across multiple fungal genera^29–32^, highlighting their potential as bioactive agents for pest control. Recent advances in metabolomics now enable systematic profiling of these metabolites, offering insights into their biosynthetic pathways and ecological roles^33–35^.

This study integrates multi-omics and morphological approaches to evaluate the biocontrol potential of *C. echinulata* isolate SMH-1, an indigenous strain isolated from a collapsed RIFA nest. We (1) quantify dose-and temperature-dependent virulence against RIFA worker castes, (2) characterize infection dynamics using scanning electron microscopy (SEM) and light microscopy, and (3) employ UPLC-MS metabolomics to identify bioactive compounds in *C. echinulata* cultures and RIFA hemolymph post-exposure. Six candidate metabolites were further assayed for insecticidal activity. Our findings establish *C. echinulata* as a promising mycopesticide candidate while elucidating its dual-mode pathogenicity, thereby advancing the development of targeted biocontrol tools against this globally invasive pest.

## 2. Results

### 3.1 Phylogenetic analysis and identification of isolate SMH-1

Phylogenetic reconstruction (Maximum Likelihood, 1,000 bootstrap replicates) positioned *Cunninghamella echinulata* isolate SMH-1. Sequence of isolate SMH-1 shown in red colour with assession number (for nucleotide sequence: SUB14919727 BSND_A05 7121) obtained from NCBI, while other groups sequences from GenBank shown in black colour. Isolate SMH-1 shared 98% cluster similarity with sequences of GenBank (*C. echinulata* group) which indicates a clear and well resolved classification (fig. 1a). Morphological characterization revealed rapid PDA colonization (Fig. 1b), producing white, densely branched mycelia with erect conidiophores (Fig. 1c). Mature conidia exhibited yellow-brown pigmentation, spinose ornamentation (Fig. 1d–f), and unispored sporangia (Fig. 1g), consistent with *C. echinulata*’s diagnostic features.

**Fig. 1:**
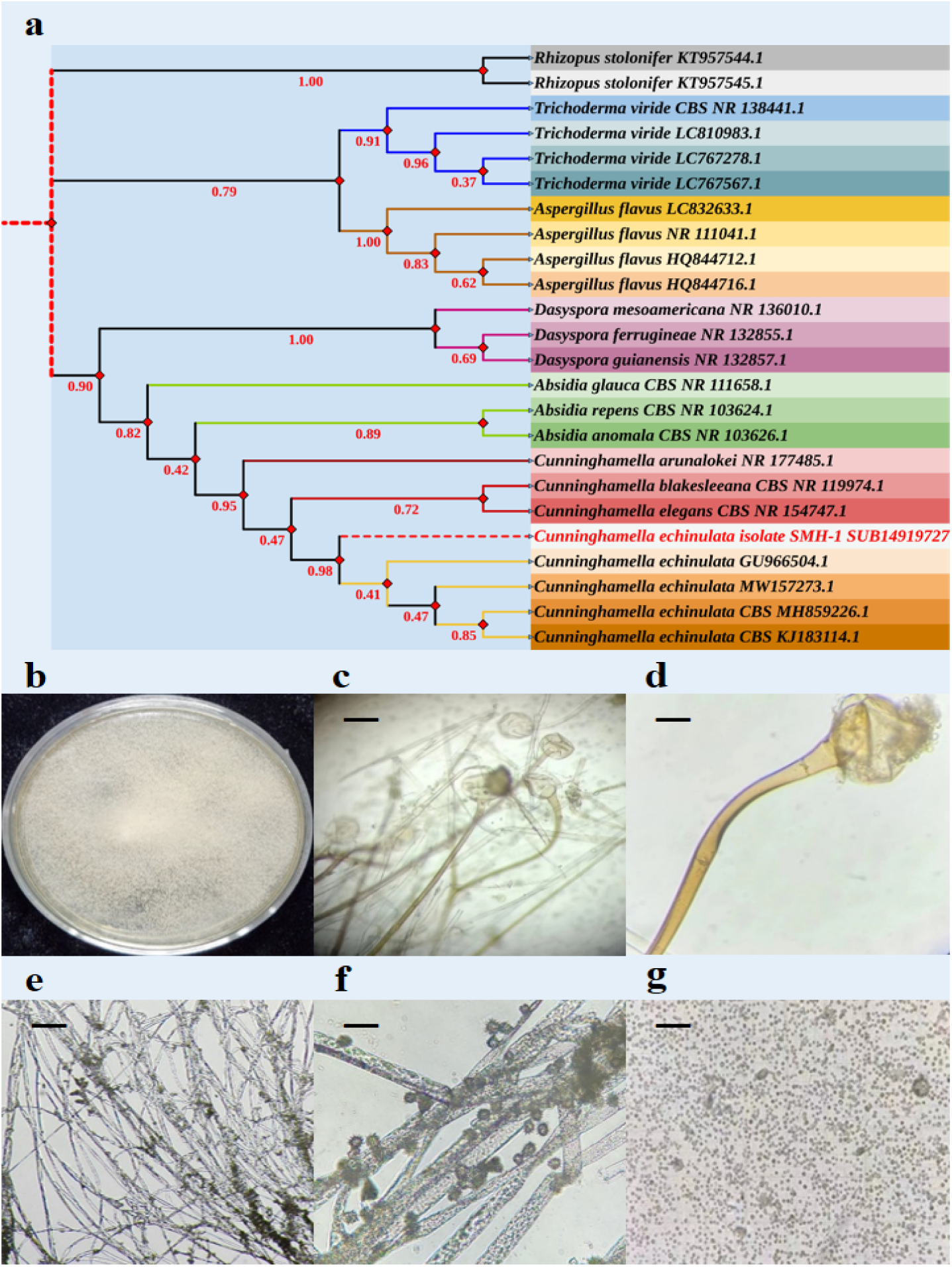
Morphological and molecular identification of the fungus *Cunninghamella echinulata* isolate SMH-1, preliminary identification was done by Dr. Deqiang Qin (Photos were taken by Dr. D.Q. Qin and Hussain Mehboob, Locality: Yunnan Agricultural University, Kunming, P.R. China); (a) Phylogenetic tree of *Cunninghamella echinulata* isolate SMH-1 constructed by the neighborhood joining method based on sample cultured from the mycelium derived from destructed nest of *Solenopsis invicta* (b) The colonial morphology cultured on potato dextrose agar (PDA) medium at day 4 (c) Mycelial morphology, (the bar indicates the 10 μm) (d) Morphology of the conidiophore, (the bar indicates the 40 μm) (e, f) characteristics of phialides, (the bar indicates the 10 μm and 40 μm, respectively) (g) morphology of the released conidia, (the bar indicates the 10 μm)

### 3.2 Pathogenicity test of isolate SMH-1 to S. invicta workers

Fungal pathogenicity was assessed by measuring the susceptibility of *S. invicta* workers (Fig. 2a). The four different concentrations of isolate SMH-1 conidial suspensions were 1×10^8^, 1×10^7^, 1×10^6^, and 1×10^5^ conidia/mL. The fig. 2a showed dose dependent mortality of *S. invicta* workers succumbed to the infection. Highest conidial suspension (1×10^8^ conidia/mL) showed mortality from day 1 while lowest conidial suspension (1×10^5^ conidia/mL) showed mortality from day 3. More than 50% mortality was found at day 5 by the highest conidial suspension (1×10^8^ conidia/mL). At high dose (1×10^8^ conidia/mL), 100% mortality of *S. invicta* workers was recorded at day 7 followed by 81.66% (1×10^7^ conidia/mL), 66.66% (1×10^6^ conidia/mL) and 65% (1×10^5^ conidia/mL). While only 3.33% mortality was found at day 7 among *S. invicta* workers treated with 0.05% Tween-80 solution as control. On the basis of this data, an approximate mean lethal concentration for 50% mortality (LC_50_) of *S. invicta* workers at day 7 calculated as 4.13×10^4^ conidia/mL.

**Fig. 2:**
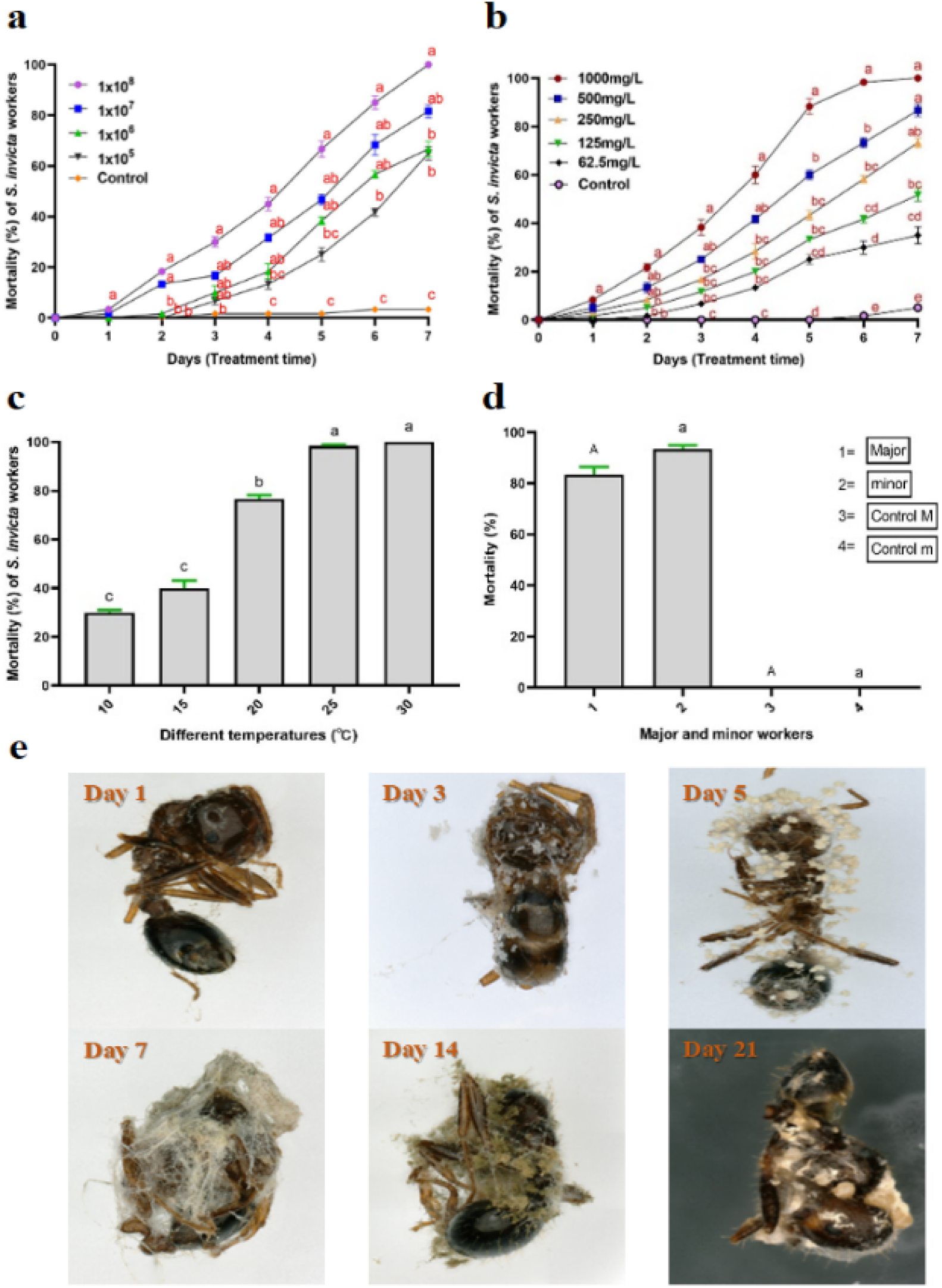
Insecticidal activity of *Cunninghamella echinulata* isolate SMH-1 against *Solenopsis invicta*; (a) Mortality rates of *Solenopsis invicta* mixed aged workers against infection by isolate SMH-1 using indicated concentrations of conidial suspensions as treatments over seven days, (b) Mortality percentage of *Solenopsis invicta* individuals after feeding 10% sugar water containing different concentrations of acetone extracts from isolate SMH-1 over 7-days, (c) Mortality percentage of workers of *Solenopsis invicta* after exposure to (5.35 × 10^7^ conidia/mL) conidial suspension indicating impact of different temperatures on isolate SMH-1’s virulence. Mortality was measured at day 7 after treatment, (d) Mortality rate of major and minor workers of *Solenopsis invicta* after exposure to (1.03 × 10^7^ conidia/mL) conidial suspension, (e) Treated workers of *Solenopsis invicta* showing the growth of *Cunninghamella echinulata* isolate SMH-1 after death. The changes in Cadaver’s of red fire ant shown over 21 days (× 200 magnification, varied light effects). Note: All experiments were performed in triplicate and each replication used 20 workers. Error bars = SD (Standard deviation), while different letters at each observation day indicate significant differences among treatments at p < 0.05 level based on Tukey’s HSD test (n = 3).

### 2.3 Toxicity of extracted metabolites to S. invicta workers

A feeding bioassay showed that acetone extracts of isolate SMH-1 possess activities against RIFA workers (see fig. 2b). The figure showed dose dependent mortality trend of RIFA workers over time with significant differences. The higher concentration (1000 mg/L) showed 100% mortality over seven days, followed by 500 mg/L, 250 mg/L, 125 mg/L and 62.5 mg/L concentrations for 86.66%, 73.33%, 51.66% and 35% mortality respectively.

### 2.4 Effect of different temperatures on ant bioassay

Effect of different temperatures on the virulence of isolate SMH-1 to *S. invicta* workers measured at day 7 by percentage mortality at dose 5.35×10^7^ conidia/mL (Fig. 2c). All different temperature treatments showed mortality from day 1, however, different temperatures showed significant differences in the mortality rate after day 7. Mortality significantly reduced to 30%, 40% for 10℃ and 15℃ temperature, respectively at day 7. At highest temperature (30℃), 100% mortality observed at same day followed by 25℃, 25℃ and 20℃ temperatures showed mortality as 98.33%, and 76.66%, respectively.

### 2.5 Virulence of isolate SMH-1 to major and minor S. invicta workers

Minor workers (<4 mm) exhibited higher susceptibility (93.3 ± 2.9%) than majors (>4 mm; 83.3 ± 3.3%) at 1.03×10⁷ conidia/mL (Fig. 2d). However, Kaplan-Meier survival analysis revealed no significant caste-specific differences (log-rank test: χ² = 2.1, p = 0.15), suggesting cuticular penetration efficiency transcends allometric scaling in polymorphic ants.

### 2.6 Median lethal concentrations of isolate SMH-1against S. invicta workers

The median lethal concentration (LC_50_) values of isolate SMH-1 against *S. invicta* workers have been evaluated with respect to different times (Table 1). Five conidial suspensions (1×10^8^, 1×10^7^, 1×10^6^, 1×10^5^ and 1×10^4^ conidia/mL) have been used. After 48 h of exposure, the median lethal concentration (LC_50_) against mixed sized *S. invicta* workers was calculated as 3.17×10^12^ conidia/mL. The lowest median lethal concentration (LC_50_) of isolate SMH-1 against *S. invicta* workers was calculated as 1×10^3^ conidia/mL at day 7 after 168h of exposure, which lead by 144 h, 120 h, 96 h, and 72 h of exposure as 1.18×10^4^, 4.83×10^4^, 4.74×10^6^, and 4.87×10^9^ conidia/mL, respectively. On the basis of median lethal concentrations of isolate SMH-1’s pathogenicity to *S. invicta* workers, no significant differences have been found among different times of exposure.

**Table 1.**
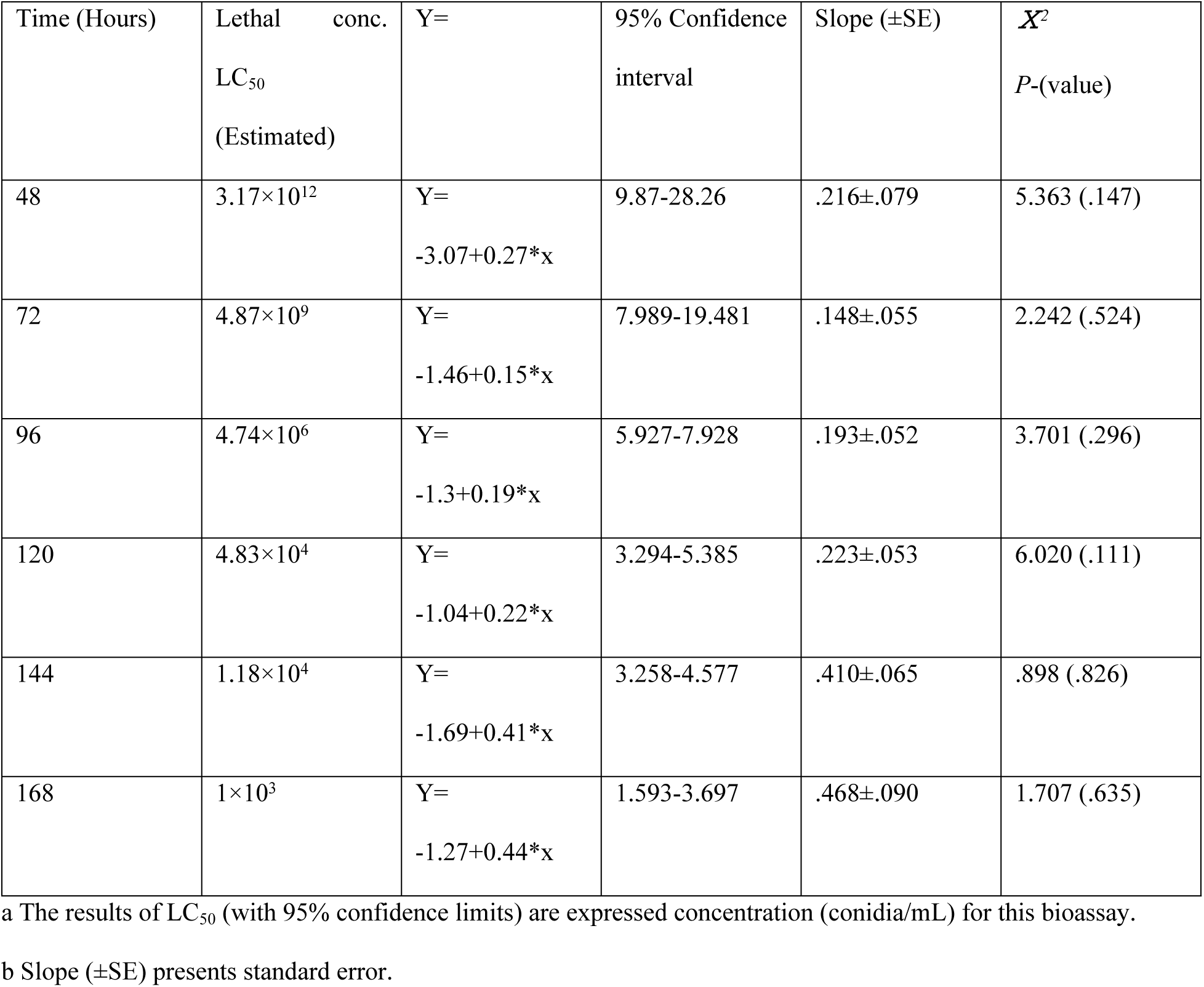
Estimated medium lethal concentration (LC_50_) of five different dilutions of isolate SMH-1 against *S. invicta* workers over seven days exposure from 48h to 168h, obtained from probit analysis (df=5). The chi-square test (*Ⅹ^2^*) value indicates the goodness of fit test at *P* ˃ 0.05.

| Time (Hours) | Lethal conc.<br>LC<sub>50</sub><br>(Estimated) | Y= | 95% Confidence<br>interval | Slope (±SE) | $\chi^2$<br>P-(value) |
| --- | --- | --- | --- | --- | --- |
| 48 | $3.17 \times 10^{12}$ | Y=<br>-3.07+0.27*x | 9.87-28.26 | .216±.079 | 5.363 (.147) |
| 72 | $4.87 \times 10^9$ | Y=<br>-1.46+0.15*x | 7.989-19.481 | .148±.055 | 2.242 (.524) |
| 96 | $4.74 \times 10^6$ | Y=<br>-1.3+0.19*x | 5.927-7.928 | .193±.052 | 3.701 (.296) |
| 120 | $4.83 \times 10^4$ | Y=<br>-1.04+0.22*x | 3.294-5.385 | .223±.053 | 6.020 (.111) |
| 144 | $1.18 \times 10^4$ | Y=<br>-1.69+0.41*x | 3.258-4.577 | .410±.065 | .898 (.826) |
| 168 | $1 \times 10^3$ | Y=<br>-1.27+0.44*x | 1.593-3.697 | .468±.090 | 1.707 (.635) |

a The results of LC<sub>50</sub> (with 95% confidence limits) are expressed concentration (conidia/mL) for this bioassay.
b Slope (±SE) presents standard error.

### 2.7 Scanned electron and light microscopy

The light microscopy revealed the infection process of isolate SMH-1 to RIFA workers over 21 days (Fig. 2e). We observed that fungal infection remained at the same angle and evenly distrusted to the whole body of RIFA. It starts from the tarsi, head and cements to the other parts. Furthermore, we observed the hyphae, conidiophore and phialides dispersing from the infected hosts over time.

The branched hyphae with conidia fully covered the cadavers over seven days. After 21 days, half of the hyphae die over dead cadavers of RIFA.

Scanned electron micrographs (SEMs) revealed that isolate SMH-1 conidia were evenly distributed throughout the body of *S. invicta* including tarsi as well. The conidial visualization shown the shriveled penetrating conidia, cat fusion of conidia over three days found on propodium and attached to, around the setae 15 and 9 days respectively. Arraying of tear shaped conidia for hyphal formation observed on ninth day as well. Additionally, germinating conidia, and elongated spines covering conidiophore at the maturity observed as well over 15 and nine days as well (fig. 3a). Similarly, SEMs revealed the fungal mycelium that fully expands to dead cadavers of RIFA over fifteen days. Mycelium found on head, thorax and abdomen on 3 and 9 days of infection respectively. Dispersing phialides and hyphae observed throughout the body of the infected host over 15 days. In addition, the mycelium mass was more in 15 days as compared to 3^rd^ day infected cadavers (Fig. 3b).

**Fig. 3:**
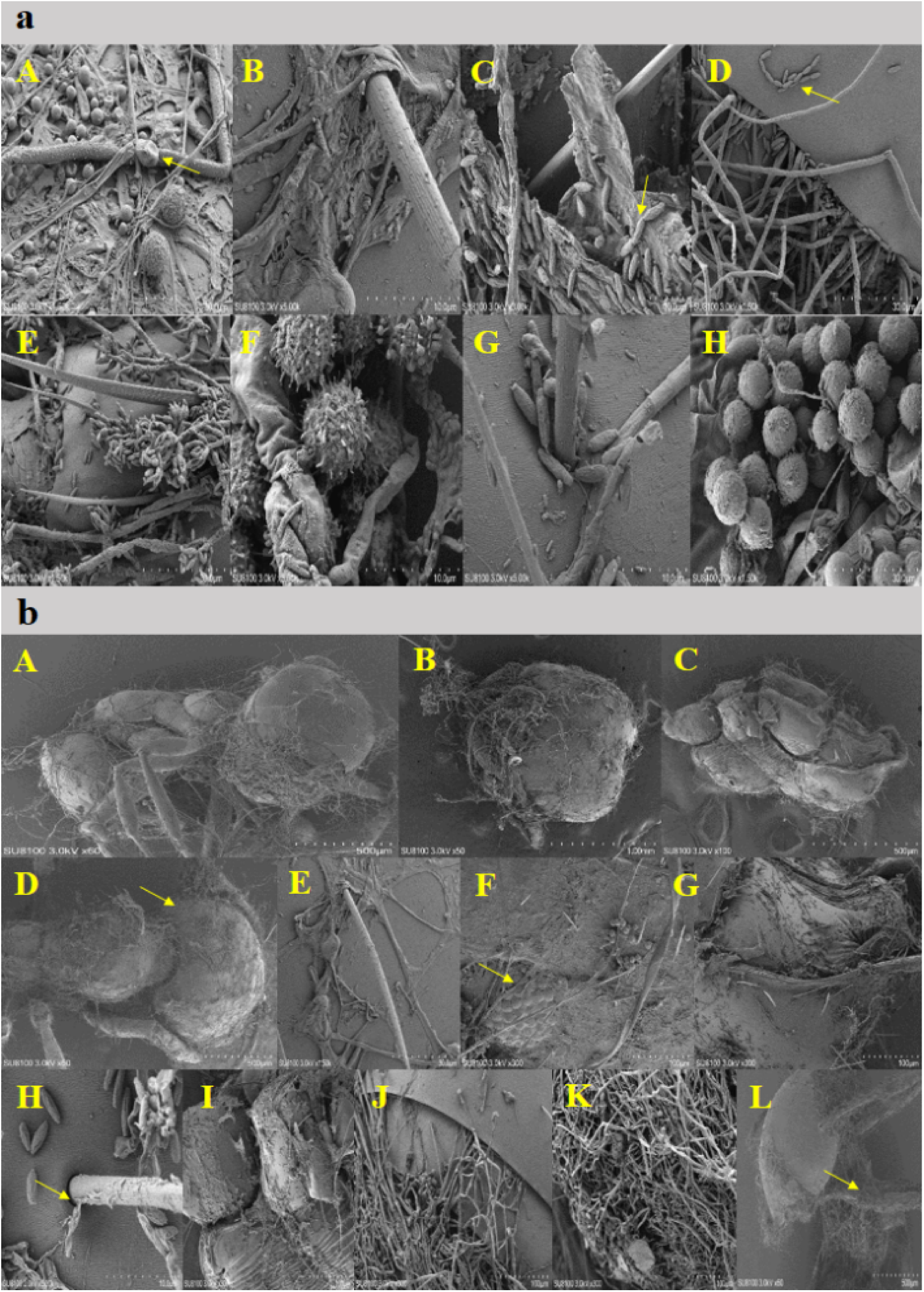
(a) Scanned electron micrographs (SEMs) revealing the fungal infection process of *Cunninghamella echinulata* isolate SMH-1 to workers of *Solenopsis invicta* over fifteen days. The fungal conidia visualized: (A) shriveled penetrating conidia, 3-day (B) attached to the setae, 15-day (C) cat fusion of conidia, 3-day (D) arraying of tear shaped conidia on the intersegmental folds in the abdomen, 9-day (E) on the propodium, 3-day (F) elongate spines covering sporangiola at maturity found on thorax grooves, 9-day (G) around the setae, 9-day (H) germinating conidia on the abdomen, 15-day (b) The fungal infection process revealed by the representatives scanned electron micrographs (SEMs). Isolate SMH-1 mycelium found: (A) on dead cadaver almost fully covered after three days of infection (B) attached to head, 3-day (C) outgrowth on the thorax, 3-day (D) on the abdomen, 9-day (E) phialides around the setae, 9-day (F) around the compound eye, 15-day (G) on the grooves of thorax, 3-day (H) at the setae, 9-day (I) between insect cuticlar folds, 15-day (J) on and between intersegmental folds of abdomen, 3-day (K) covering the tip of abdomen, 15-day (L) on the surface of legs, 15-day.

#### 2.8.1 Metabolic discriminant analysis of isolate SMH-1

Samples were analyzed by multivariate statistical principal component analysis (PCA) (Fig. 4a). Among the three principal components, the contribution rate of principal component 1 was 60%, and the contribution rate of principal component 2 was 18.03%. The cumulative interpretation rate of A vs B in the X-axis direction is R2X=0.743, R2X=0.939 for A vs C, and R2X=0.929 for B vs C, so the PCA model has a good fit.

**Fig. 4:**
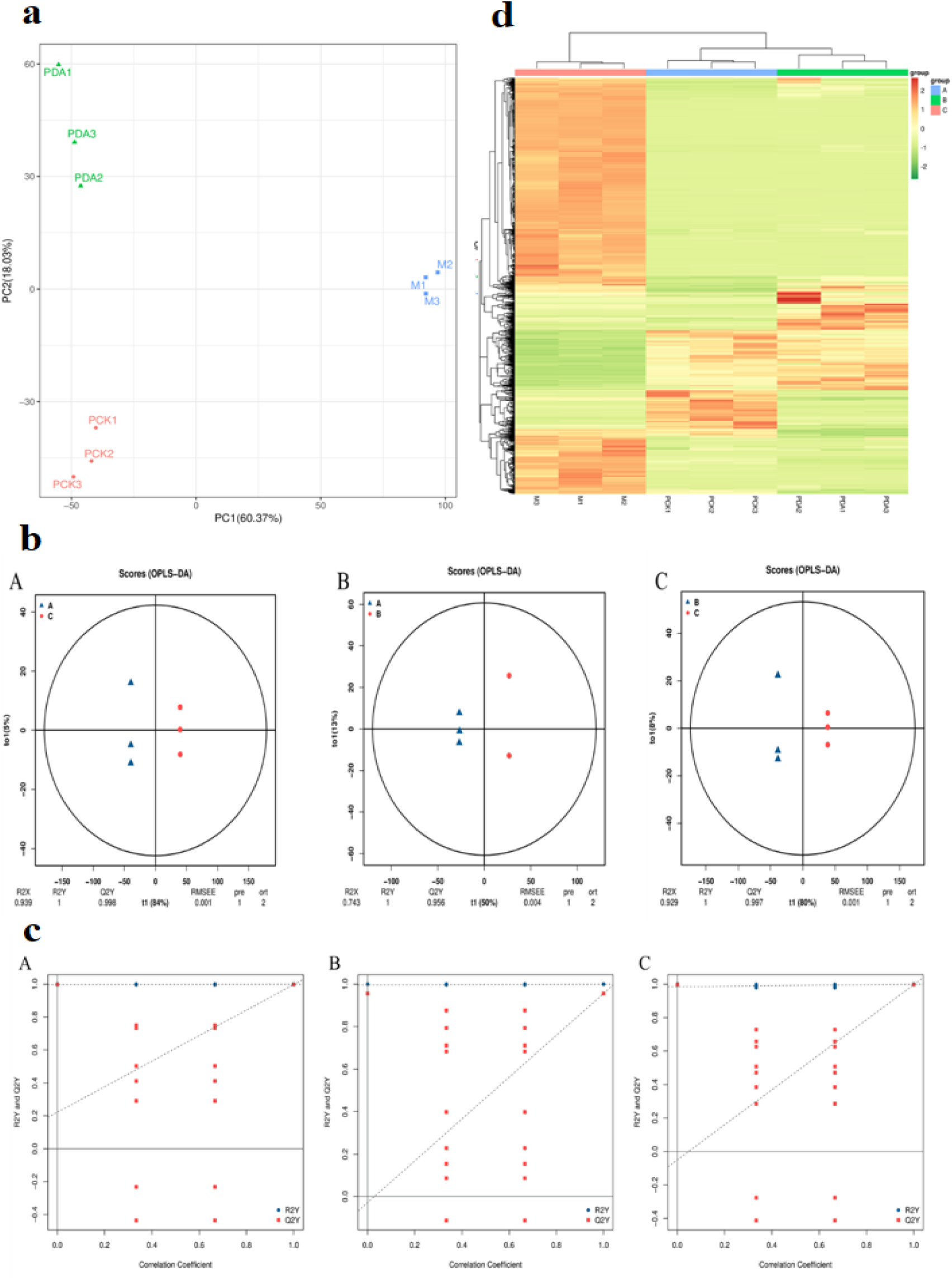
UPLC-MS based metabolic profiling of *Cunninghamella echinulata* isolate SMH-1; (a) PCA map of metabolic profiles in isolate SMH-1 from blank PDA medium (A), the PDA medium cultured with hyphae (B) and the mycelium (C), (b) OPLS-DA score plot of metabolic profiles in isolate SMH-1 (A: PCK vs M; B:PCK vs PDA; C:PDA vs M), (c) OPLS-DA permutation test plots for each group (A: PCK vs M; B:PCK vs PDA; C:PDA vs M), (d) Cluster heat map of each group of samples from isolate SMH-1.

#### 2.8.2 Orthogonal Partial Least Squares Discriminant Analysis (OPLS-DA)

In order to understand the differences between the fungi themselves and the secreted metabolites, the differences between the blank PDA medium (A), the PDA medium cultured with hyphae (B) and the mycelium compounds (C) were analyzed separately, and the orthogonal partial least squares discriminant analysis (OPLS-DA) model was used to test the differences. OPLS-DA analysis is a multivariate statistical analysis method with supervised pattern recognition that effectively excludes non-study-related effects to screen for differential metabolites. The prediction parameters of the evaluation model are R2X, R2Y and Q2, where R2X and R2Y represent the interpretation rate of the model to X and Y matrices respectively, Q2 represents the prediction ability of the model, the closer these three indicators are to 1, the more stable and reliable the model is, Q2>0.5 is considered to be an effective model, and Q2 is >0.9 is an excellent model. The values of R2X, R2Y and Q2 in the three groups are all close to 1, indicating that the evaluation model is reliable and valid. OPLS-DA was used to analyze groups A, B, and C in pairs to plot score plots and model displacement test charts (fig. 4b,c), and the score plots and model permutation test charts of OPLS-DA showed that there was significant separation in the different comparison groups.

#### 2.8.3 Differential metabolite analysis

Based on metabolome data analysis, we found 812 compounds, classified according to their compound category, we divided the differential compounds into 17 categories of compounds, which are monosaccharides, oligosaccharides, neurotransmitters, other hormones, steroid hormones, fatty acids, phospholipids, bases, nucleosides, nucleotides, carboxylic acids, amines, amino acids, peptides, 27-carbon atoms, cofactors, and vitamins, among which the most differentiated compounds are phospholipids, the number of which is 11; the differential compounds were LysoPC (20:5 (5Z, 8 Z, 11Z, 14 Z, 17Z)); LysoPC (20:4(5Z,8Z,11Z,14Z)); PG (i-12:0/a-17:0); LysoPC (18:1(11Z)); PE (15:0/16:1(9Z)); PC (P-18:0/0:0); LysoPC (18:0); PS (16:0/18:2(9Z,12Z)); LysoPC (18:3(6Z,9Z,12Z)); LysoPC (16:1(9Z)/0:0); LysoPC (16:0); the least different compounds were steroid hormones and other hormones, with a number of 1; The differential compounds were histamine and calcitriol, respectively. We classify the compounds according to the different metabolic pathways involved in them, and there are 17 differential metabolic pathways. Transport and catabolism, membrane transport, signal transduction, signaling molecules and interaction, folding, sorting and degradation, translation, amino acid metabolism, biosynthesis of other secondary metabolites, carbohydrate metabolism, energy metabolism, glycan biosynthesis and metabolism, lipid metabolism, metabolism of cofactors and vitamins, metabolism of other amino acids, nucleotide metabolism, xenobiotics biodegradation and metabolism, sensory system; among them, 73 compounds were found in the amino acid metabolism pathway, mainly including caldine, spermine, dihydroxyphenylacetic acid etc.; among them, there was one compound involved in the folding, sorting and degradation metabolic pathway, which was S-adenosylmethionine.

In order to more visually demonstrate the relationship between samples and the differences in the expression of metabolites between different samples, we performed a heat map analysis of the expression of significantly differentially differentiated metabolites enriched in the pathway (see Fig. 4d). Green is the low-expressing substance, and orange is the high-expressing metabolite. Group C was significantly separated from groups A and B, indicating that there were significant differences between group C, groups A and B, and the significantly different metabolites screened could be used as markers to distinguish the three groups. Metabolites clustered together have similar functions or participate in the same metabolic pathway.

#### 2.8.4 Analysis of significantly differential metabolites

Combined with the P-value value, the metabolites with significant differences between different groups were further screened out, and the metabolites with P-value<0.05 were considered to be significantly different. As shown in Fig. 5a, there were 560 metabolites (186 up-regulated and 374 down-regulated) in group A and group B, 1668 metabolites (1294 up-regulated and 374 down-regulated) in group A and group C, and 1575 metabolites (1250 up-regulated and 325 down-regulated) in group B and group C.

**Fig. 5:**
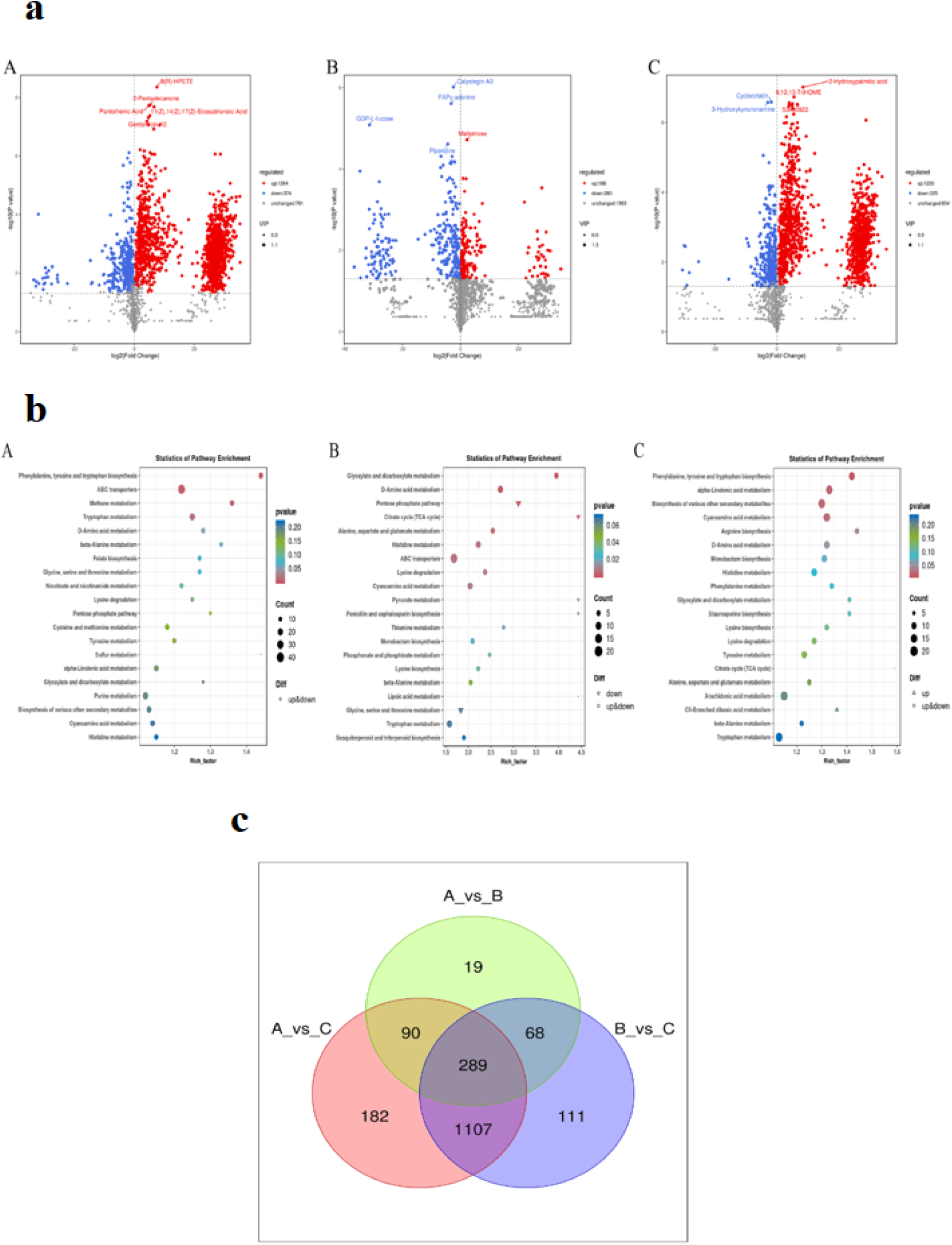
(a) Volcanic map of differential metabolites from isolate SMH-1 samples (A:PCK vs M; B:PCK vs PDA; C:PDA vs M), (b) KEGG enrichment of differential metabolites (A:PCK vs M; B:PCK vs PDA; C:PDA vs M), (c) Venn diagram of unique differential metabolites between different groups.

#### 2.8.5 Analysis of differential metabolite metabolism pathways

All the differential metabolites in the different comparison groups were matched to KEGG’s database to obtain information on the pathways in which the metabolites were involved. The annotated results were enriched to obtain pathways with more differential metabolite enrichment, as shown in Fig. 5b. Differential metabolites are mainly annotated and enriched in tryptophan metabolism, methane metabolism, glyoxylic acid and dicarboxylic acid metabolism, D-A amino acid metabolism, pentose phosphate pathway, citric acid cycle, biosynthesis of various other secondary metabolites, and cyanamide metabolism.

#### 2.8.6 Analysis of specific differential metabolites

The Venn diagram shows that there are 19 unique differential metabolites in A vs B, 182 unique differential metabolites in A vs C, and 111 unique differential metabolites in B vs C (Fig. 5c).

#### 2.8.7 Toxicity of six different compounds to S. invicta workers

The feeding bioassays showed that different concentrations of fungal compounds possess activities against RIFA workers (see Fig. 6). The figure showed dose dependent mortality trend of RIFA workers over time with significant differences. The highest concentration (100mg/l) showed 100% mortality of worker ants over 12 days for dihydrocoumarin (Fig. 6d), followed by coumarin (Fig. 6c), 5-methoxysporalen (Fig. 6a), 7-hydroxycumarin (Fig. 6b), scopoletin (Fig. 6e) and scopolin (Fig. 6f), over 14 and 15 days respectively. The lowest concentration (1 mg/L) showed highest mortality of 18.33% for 7-hydroxycumarin over 15 days followed by dihydrocoumarin, 5-methoxysporalen, scopoletin, scopolin and coumarin as 16.66%, 15%, 15%, 11.66% and 8.33% respectively.

**Fig. 6:**
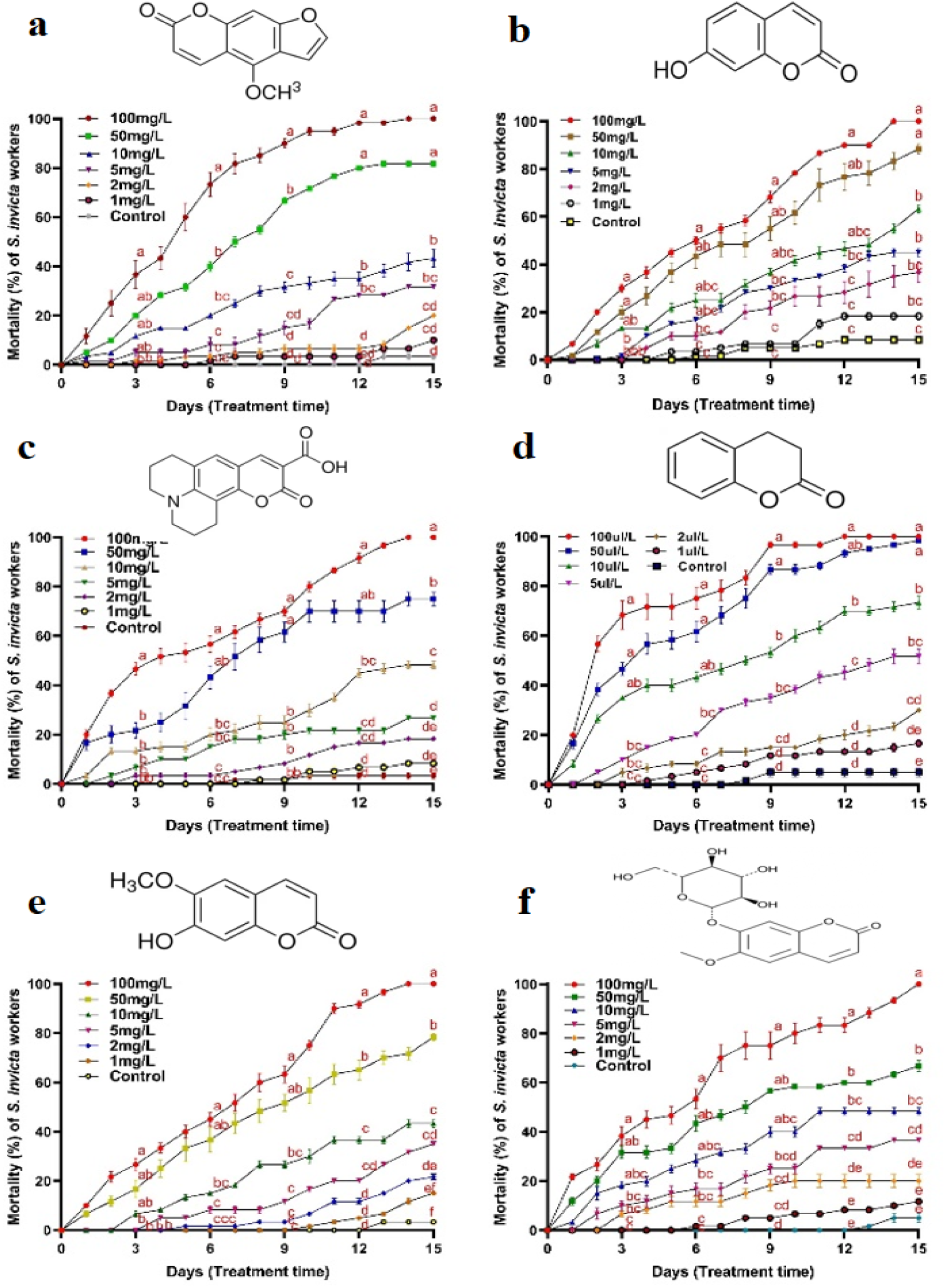
Mortality percentage of *Solenopsis invicta* individuals after feeding 10% sugar water containing different concentrations of six compounds over 15 days; compounds are (a) 5-methoxysporalen, (b) 7-hydroxycoumarin, (c) coumarin, (d) dihydrocoumarin, (e) scopoletin, (f) scopolin. Error bars showing the standard deviation, while all experiments were performed in triplicate. Different letters at each observation day indicate significant differences among treatments at p < 0.05 level based on Tukey’s HSD test (n = 3).

#### 2.9.1 Analysis of differentially expressed metabolites in RIFA

Primarily, a general impression of the clustering information between different groups was obtained using an unsupervised principal component analysis (PCA) method without any sample designation. Based on the likeness between the metabolic profiles of samples, the groupings in PCA scores plot attained.

The PCA scores plot showed clear separations between the compound dihydrocoumarin (A), metabolites (B), conidia (C), and control (CK) groups (see Fig. 7a). Meanwhile, the compound dihydrocoumarin (A) group was clearly separated from conidia (C) group along the second principal component (PC2), which explained 30.05% of the total variance. However, clear separations were observed between the control (CK) group and different treatment (A, B and C) groups along the both principal components, while the principal component one (PC1) and two (PC2) explained 43.54% and 30.05% of the total variance respectively.

**Fig. 7:**
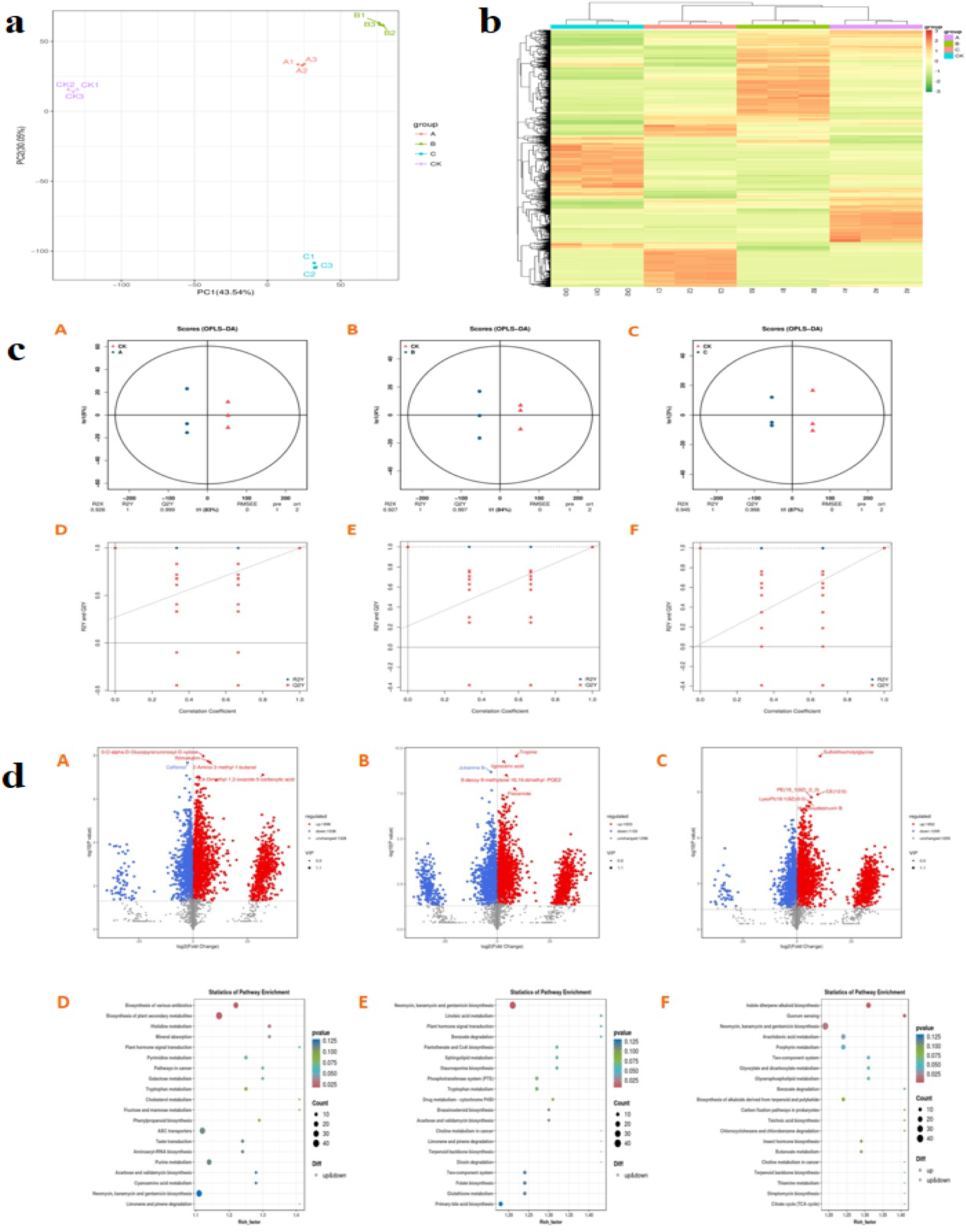
Metabolic profiling of *Solenopsis invicta* based on LC-QTOF in response to isolate SMH-1; (a) PCA score plot of metabolic profiles in *Solenopsis invicta* workers from compound dihydrocoumarin (A), metabolites (B), conidia (C) and control (CK) groups, (b) Hierarchical clustering heatmap of differential metabolite union [Note: A: Compounds (A) vs Control (CK), B: Metabolites (B) vs Control (CK), C: Conidia (C) vs Control (CK)]; (c) OPLS-DA Score Plot and OPLS-DA permutation test plots [Note: A: Compounds (A) vs Control (CK), B: Metabolites (B) vs Control (CK), C: Conidia (C) vs Control (CK); D: Compounds (A) vs Control (CK), E: Metabolites (B) vs Control (CK), F: Conidia (C) vs Control (CK)], (d) Volcanic plot of differential metabolites from isolate SMH-1 samples [Note: A: Compounds (A) vs Control (CK), B: Metabolites (B) vs Control (CK), C: Conidia (C) vs Control (CK)]; KEGG enrichment of differential metabolites in *Solenopsis invicta* [Note: D: Compounds (A) vs Control (CK), E: Metabolites (B) vs Control (CK), F: Conidia (C) vs Control (CK)].

#### 2.9.2 Potential metabolite search and identification

Like partial least squares (PLS), the orthogonal partial least squares discriminate analysis (OPLS-DA) model is a supervised method that generally provides greater discrimination power compared to PCA model. OPLS-DA models were applied between the model and other groups to improve the variation. OPLS-DA score plots and model permutation test charts showed a clear separation among the control group and different treatment groups without overlapping (Fig. 7c). This outcome of OPLS-DA models supports the above PCA score plots results. The quality of OPLS-DA models was assessed on the basis of R2X, R2Y and Q2 values. As the R2X and R2Y values were close to 1 and Q2 values were over 0.5, it showed that the models were valid and possess high predictive abilities.

To further explore the relationship between treatments and the differentially expressed metabolites, the heat map analysis performed (Fig. 7b). The low-expressing metabolites indicated as green and orange is the highly-expressing metabolites. From the Fig. 7b, we can see the group B (metabolites) is clearly separated from the groups A (dihydrocoumarin) and C (conidia). This indicated the significant differences among differentially expressed metabolites between different treatment groups A, B, and C. The significantly different metabolites screened could be used as markers to distinguish the three groups.

#### 2.9.3 Significantly differential metabolites and pathway analysis

As the volcano plot provides a quick visualization of the overall trend in metabolite abundance differences between different groups and the statistical significance of these differences. So, based on the results of OPLS-DA, differential metabolites further screened out on the basis of VIP values obtained from the multivariate analysis of the OPLS-DA model. Numerous metabolites have been significantly increased or decreased or remained unaffected in different treatment groups A, B, and C (fig. 7d). There were 2934 metabolites (1896 up-regulated and 1038 down-regulated) in group A (Compound), 2966 metabolites (1833 up-regulated and 1133 down-regulated) in group B (metabolites) and 3057 metabolites (1852 up-regulated and 1205 down-regulated) in group C (conidia). For the group A treatment, among top five metabolites 3-O-alpha-D-Glucopyranuronosyl-D-xylose, rilmakalim, 2-Amino-3-methyl-1-butanol, and 2,4-Dimethyl-1,3-oxazole-5-carboxylic acid showed marked increase while caffeinol showed the marked decrease. For the group B treatment, tropine, lignoceric acid, 9-deoxy-9-methylene-16,16-dimethyl-PGE2, flecainide showed marked increase and jubanine B showed significantly marked decrease. While for group C, top five metabolites all showed marked increase were sulfolithocholylglycine, PE (18_1(9Z)_0_0), CE (12:0), LysoPI (18:1(9Z)/0:0) and hydroxydestruxin B. These significant alterations of metabolites among different treatments and control provide insights on their effect on different pathways.

MetaboAnalyst 4.0 (Free web based tool that provides high quality KEGG metabolic pathway databases) was used to resolve all differentially expressed metabolites. To perform the pathway analysis, all the differential metabolites in the different comparison groups were matched to KEGG’s database to obtain information on their involvement in different pathways. The annotated results were enriched to obtain pathways with more differential metabolite enrichment shown above (fig. 7d). Several common and specific pathways were observed in different treatment groups. However, differential metabolites are mainly annotated and enriched in neomycin, kanamycin and gentamicin biosynthesis, two-component system, benzoate degradation, choline metabolism in cancer, terpenoid backbone biosynthesis, plant hormone signal transduction, tryptophan metabolism, acarbose and validamycin biosynthesis, limonene and pinene degradation. On the basis of both P and impact values, the key pathways observed for group C (Conidia) as Indole diterpene alkaloid biosynthesis, Quorum sensing, Arachidonic acid metabolism, Porphyrin metabolism, glyoxylate and dicarboxylate metabolism, glycerophospholipid metabolism, biosynthesis of alkaloids derived from terpenoid and polyketide, carbon fixation pathways in prokaryotes, teichoic acid metabolism, chlorocyclohexane and chlorobenzene degradation, insect hormone biosynthesis, butanoate metabolism, thiamine metabolism, streptomycin biosynthesis, citrate cycle (TCA cycle). While Linoleic acid metabolism, pantothenate and CoA biosynthesis, sphingolipid metabolism, staurosporine biosynthesis, phosphotransferase system, drug metabolism, cytochrome P450, brassinosteroid biosynthesis, dioxin degradation, folate biosynthesis, glutathione metabolism, primary bile acid biosynthesis observed as key pathways for group B (metabolites). Similarly, the only key paths for group A (compound) observed as biosynthesis of various antibiotics, biosynthesis of plant secondary metabolites, histidine metabolism, mineral absorption, pyrimidine metabolism, pathways in cancer, galactose metabolism, cholesterol metabolism, fructose and mannose metabolism, phenylpropanoid biosynthesis, ABC transporters, taste transduction, aminoacyl-tRNA biosynthesis, purine metabolism, cyanoamino acid metabolism.

The figure shown that there are 4 differentially regulated metabolites for group C (conidia) treatment which could reflect the main differentially metabolic response of RIFA (Fig. 8a). As shown in (Fig. 8b), 3 differentially regulated metabolites for group B (metabolites) could reflect the differential metabolic response of RIFA. Similarly, 2 differentially regulated metabolites can be seen for group C (dihydrocoumarin) treatement (Fig. 8c). It could draw a whole overview of differential metabolic changes of RIFA to C, B and A treatments and useful to understand their killing mechanisms respectively. Supplementary table 2 presents the values of differentially expressed metabolites impacting the energy production mechanism, citrate cycle (TCA cycle) of *S. invicta*. The levels of pyruvate, pyruvic acid and succinic acid were lower in workers of *S. invicta* after seven days of treatment as compared to control, while the fumaric acid was higher than control group after seven days of treatment with conidia. After seven days of feeding with fungal acetone extratcs, the level of pyruvate was higher in treated workers as compared to control, while the levels of pyruvic acid and succinic acid were lower. On the other hand, the levels of both pyruvic acid and fumaric acid were lower after seven days of feeding with compound dihydrocoumarin.

**Fig. 8:**
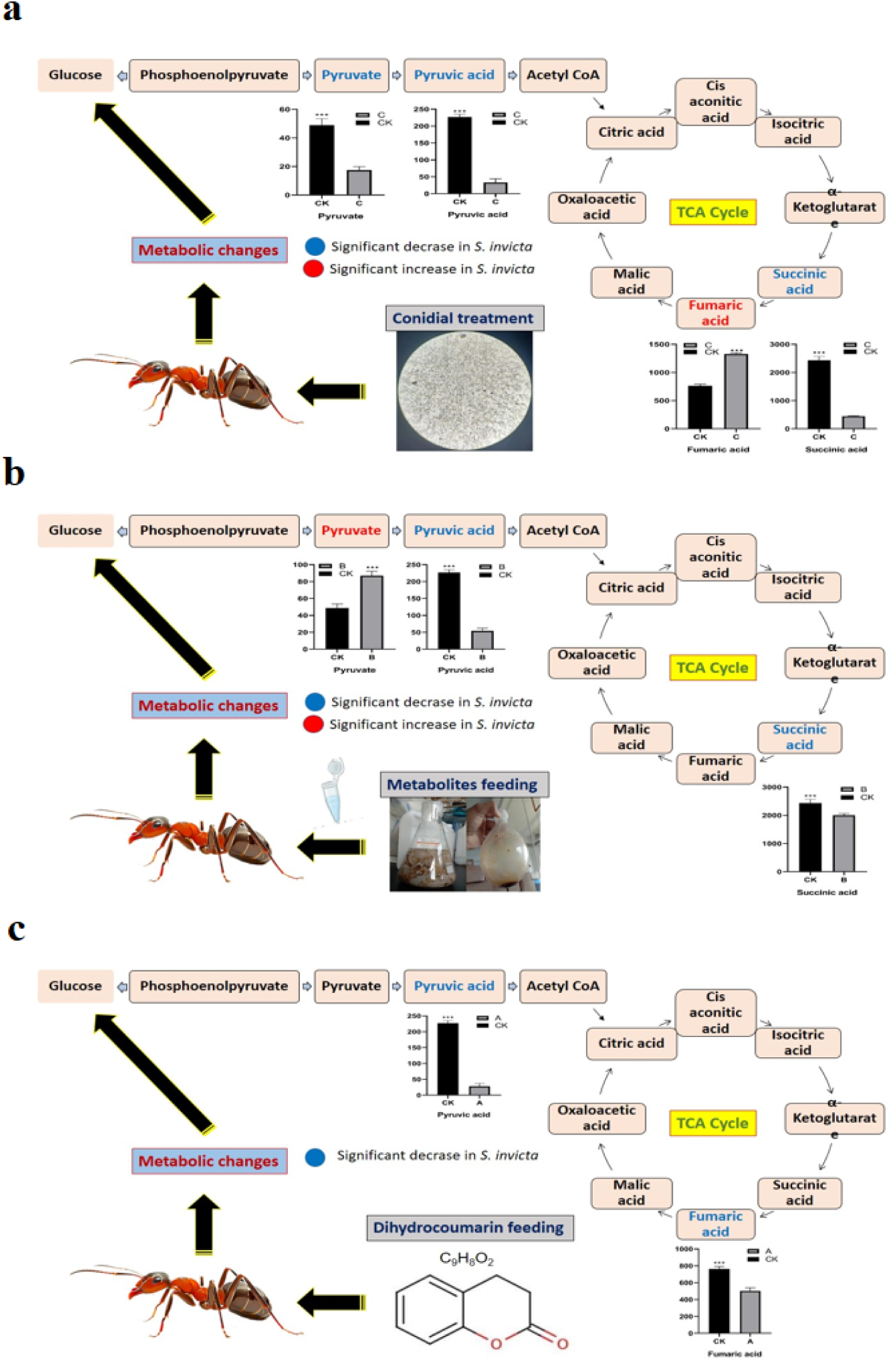
Schematic impression of primarily affected metabolic pathway of *S. invicta* due to *Cunninghamella echinulata* isolate SMH-1; (a) conidial treatment, (b) acetone extracts feeding, (c) dihydrocoumarin feeding; *** indicates the metabolite in this group is the differentially regulated metabolite (VIP > 1 & P < 0.05) screened out from the results of multivariate and univariate analysis as shown in Fig. 7.

## 3. Discussion

Prior studies have established the efficacy of entomopathogenic fungi such as *B. bassiana* and *M. anisopliae* against *S. invicta*^16–20^. Our findings extend this paradigm by demonstrating *C. echinulata* isolate SMH-1’s potent virulence, achieving 100% mortality in RIFA workers at 1×10⁸ conidia/mL within 7 days. This aligns with reports of *C. echinulata*’s pathogenicity against *Galleria mellonella*^46^, *Acytopeus curvirostris persicus*^47^, and *Culex pipiens*^48^, underscoring its broad-spectrum entomopathogenic potential. The observed cuticular penetration and systemic colonization corroborate established infection mechanisms involving hydrophobic conidial adhesion and enzymatic degradation^49^, positioning SMH-1 as a viable biocontrol agent.

Bioassay revealed the influence of different temperatures on the virulence of isolate SMH-1 to *S. invicta* workers. For the dose, 5.35×10^7^ conidia/mL, mortality trend observed over seven days. While mortality was significantly reduced to 30% and 40% for 10℃ and 15℃ temperature, respectively. Further, the virulence of isolate SMH-1 against major and minor workers was assessed at concentration 1.03×10^7^ conidia/ml; after 7 days, minor workers showed highest mortality as 93.33% followed by the major workers as 83.33%. Park et al. (2022) reported the 100% mortality of major and minor workers of *S. invicta* after 7 and 8 days of exposure to 1×10^7^ conidia/mL concentration of *B. bassiana* ANU1. This study also revealed the decrease in virulence of fungus at low temperatures. Mortality reduced to 26.6% and 20% for major and minor workers, respectively^40^. Generally, the optimal temperatures for germination, sporulation, growth, and virulence of entomopathogenic fungi have been reported between 20-30℃^50,51^, while temperature may importantly marked environmental factor affecting the survival and capability to cause the host mortality^51–53^. Because the RIFA has capability of foraging between 15-43℃^54^, so, even at the low temperature, the mortality showed good infectivity potential of isolate SMH-1. This could be helpful to get more premised results in future about its activity under varied environmental conditions.

To better understand and confirm the fungal infection process, we integrated light microscopy and electron microscopy technique. For both observations, we found that fungal infection remained uniform and evenly distributed throughout the whole body of RIFA cadavers, we found the conidial attachment from tarsi to head and it’s dispersing to the other parts over course of time. Moreover, the conidial and fungal mycelium visualization through SEMs exposed the whole infection process including conidial germination, conidial penetration, cat fusion of conidia, arraying of conidia, hyphae formation, dispersing phialides and hyphae throughout the whole surface of dead cadavers. Similar findings have been reported by Hussain et al. (2021), their study demonstrated that *Cordyceps javanica* was capable to infect different regions of the host ultimately leading to increase the rate of host mortality and conidial production via colonization of the *Diaphorina citri* cadaver^37^. Zhang et al. (2021) also reported that the conidia of *B. bassiana*, strain BbYT12 have the ability to attach at any site of the host^55^. Liu et al. (2023) also found that the conidia of *Lecanicillium araneicola* strain HK-1 are also capable to bind with feet and dorsum of *Aphis craccivora*^56^. Early binding of the entomopathogenic fungal conidia to the host cuticle heavily depends on a combination different hydrophobic interactions and adherence union^57,58^. Our findings showed the adherence capability of isolate SMH-1 conidia to any site of the RIFA cadavers, though, the observance of fungal mycelium and conidia on to the setae, intersegmental folds and grooves were particular, but it was also in line with findings of Toledo et al (2010)^59^. The possible reason for frequent trapping of the conidia by setae may be due to the favorable environment of the setae, as it relatively humid because of respiration process^60,61^, which is helpful in conidial germination and fungus activity.

Earlier studies demonstrated that entomopathogenic fungi produce a wide range of fungal secondary metabolites, which are generally low molecular weight organic compounds. It is also confirmed that these secondary metabolites facilitates in fungal pathogenicity by depressing the target’s immunity, which ultimately leads to damaged nervous system and make target species more susceptible. These are chemically classified as amino acid derivatives, peptides, peptide hybrids, cyclic depsipeptides, terpenoids, and polyketides^27,28^. Our data showed that crude extracts of isolate SMH-1 retain high insecticidal potential against RIFA. Over 7 days of treatment, 100% mortality observed for 1000mg/l dose, while 86.66%, 73.33%, 51.66% and 35% mortality observed against 500 mg/L, 250mg/L, 125mg/L and 62.5mg/L concentrations, respectively. Santamarina et al. (2002) observed 100% mortality of large milkweed bug induced by dichloromethane extracts of *P. oxalicum* cultured from Wickerham broth^62^. Arunthirumeni et al. (2023) also reported the larvicidal activity of crude extracts from culture of a *Penicillium* strain related to *P. chrysogenum* caused significant mortality in *Spodoptera litura* and *Culex quinquefasciatus* populations^31^. Moreover, another study revealed the inhibited phagocytic activity of *Plutella xylostella* which ultimately lowering its resistance, due to rapid decrease in hemocytes induced by crude extracts of *Isaria fumosorosea*. After 6 days of treatment, 91 mortality observed in *P. xylostella* population^63^. Vinayaga-Moorthi et al (2015) also observed that the secondary metabolites of *I. fumosorosea*, *Beauveria bassiana*, and *Paecilomyces variotii* notably impacted the feeding, growth, fecundity, and hatchability of *S. litura*. This suggested the potential of crude extracts to be used in management of target-insect pest^64^.

Mass spectrometry (MS) technique has become a popular choice by scientists for a wide range of metabolic profiling studies. MS is capable of deriving a wide dynamic range of metabolites, compounds detection in extremely complex molecules and reproducibility of quantitative analysis^65^. Ultra-performance liquid chromatography (UPLC) coupled with triple quadmass spectrometry is profound method for the detection of metabolites, metabolites profiling and detection of active (chemical) compounds^66^. Our findings explored the differences in metabolites between the blank PDA medium, the PDA medium cultured with hyphae, and the mycelium itself using UPLC-MS analysis. In total we found 812 metabolites and chemically classified those according to the particular category. Seventeen categories of differential metabolites were found as monosaccharides, oligosaccharides, neurotransmitters, other hormones, steroid hormones, fatty acids, phospholipids, bases, nucleosides, nucleotides, carboxylic acids, amines, amino acids, Peptides, 27-Carbon atoms, cofactors, and vitamins. Significantly differentiated eleven metabolites were from phospholipids category. Furthermore, Venn diagram showed 19 unique differential metabolites between blank PDA medium and PDA medium cultured with mycelium, 182 unique differential metabolites in blank PDA medium and mycelium, and 111 unique differential metabolites between PDA medium cultured with mycelium and mycelium. Moreover, KEGG database revealed the relevant pathway information, in which metabolites were mainly involved and annotated. The main pathways found were tryptophan metabolism, methane metabolism, glyoxylic acid and dicarboxylic acid metabolism, D-A amino acid metabolism, pentose phosphate pathway, citric acid cycle, biosynthesis of various other secondary metabolites, and cyanamide metabolism.

Yang et al. (2023) found 74 common metabolites in normal and degenerated strains of *M. anisopliae* based on gas chromatography–mass spectrometry (GC–MS) metabolomics approach.

The 40 metabolites of which were significantly differentiated. Additionally, KEGG enrichment analysis identified 47 significant pathways of which mainly annotated pathways were alanine, aspartate and glutamate metabolic pathways and the glycine, serine and threonine metabolism^67^. Similarly, Bhadani et al. (2021) found 105 metabolites common in five strains of *B. bassiana* through metabolomics approach, volcano plot analysis revealed 58 compounds significantly diverse between JAU1 and JAU2 strain^30^. Wang et al. (2024) also reported the differentially expressed metabolites between mycelium and fermentation broth of *Penicillium restrictum*, total of 799 intracellular and 468 extracellular differential metabolites found. Further, KEGG analysis revealed that mainly annotated and highly enriched extracellular pathways were alanine, aspartate and glutamate metabolism, glyoxylate and dicarboxylate metabolism, and terpenoid backbone biosynthesis^68^. This established strong basis of metabolomics approach in revealing secondary metabolites. On this basis, different compounds can be harvested according to their functioning.

Recently, the use of entomopathogenic fungi garnered significant attention from the experts to be used in management of different insect pests. However, their main application rely on the conidial treatment which has been reported to be environmentally sensitive. So, alternatively several biopesticides have been formulated based on secondary metabolites derived from entomopathogenic fungus. These active compounds or biomolecules offer great benefit as biocontrol agent and could be categorized as cyclic depsipeptides, amino acids, polyketides, polyphenols and terpenoids etc. Secondary metabolites use in agricultural pest management opens the more efficient and safe way of application. However, their secondary effects, type, nature and delivery methods should be considerate too^69^. In this study, based on the metabolomics profiling of isolate SMH-1, we tested the activity of six different compounds selected/derived from isolate SMH-1. The feeding bioassays revealed their potential against RIFA workers. The active compound dihydrocoumarin showed 100% mortality at 100mg/l concentration after 12 days of treatment. Coumarin, 5-methoxysporalen, 7-hydroxycumarin, scopoletin and scopolin also showed 100% mortality of treated RIFA at 100mg/l concentration, after 14 and 15 days, respectively. While the lowest concentration (1mg/l) showed highest mortality of 18.33% for 7- hydroxycumarin over 15 days followed by dihydrocoumarin, 5-methoxysporalen, scopoletin, scopolin and coumarin as 16.66%, 15%, 15%, 11.66% and 8.33% respectively.

Several studies reported the efficacy of coumarins and their derivatives against insect pests^70–74^. Xia et al. (2023) reported significant reduction in the growth and development of *S. litura* after exposure with 1%, 2% and 3% coumarin through diet^75^. Vargas-Soto et al. (2017) also demonstrated that coumarin derivatives possess strong activity against *Drosophila melanogaster* larvae and reported 30-80% mortality between 10-100 µg/mL concentrations^76^. Similarly, Liu et al. (2023) synthesized 28 new derivatives of scopoletin and tested their activity against phytophagous mite *Tetranychus cinnabarinus* and the brine shrimp *Artemia salina*, the compounds 5a and 5j found promising with LC_50_ values as 57 and 20 μg/mL against *T. cinnabarinus*^77^. Our findings are consistent with these studies. It also suggested the potential of target compounds against RIFA and these compounds could be used as alternative to chemical pesticides in RIFA management. In addition, it also opens new ways to search for entomopathogenic fungal secondary metabolites with novel structures and insecticidal potential.

The sound response of an insect to a pathogenic infection exposes by significant alterations in its primary and secondary metabolites^78,79^ (Li et al. 2020; Alfaro et al. 2021). In this study, we used LC-QTOF based metabolomics approach to demonstrate the differentially expressed metabolites of RIFA in response to isolate SMH-1 conidia, its acetone extracts and dihydrocoumarin compound treatment. Additionally, to draw an overall disturbed metabolic response of RIFA against particular treatment. We observed that for the conidial treatment top five metabolites showed significant up regulation were sulfolithocholylglycine, PE(18_1(9Z)_0_0), CE(12:0), LysoPI(18:1(9Z)/0:0) and hydroxydestruxin B. Among top five metabolites of fungal extract treatment, tropine, lignoceric acid, 9-deoxy-9-methylene-16,16-dimethyl-PGE2, and flecainide showed marked up-regulation, while jubanine B showed significant down-regulation. Similarly, for the dihydrocoumarin compound, 3-O-alpha-D-Glucopyranuronosyl-D-xylose, rilmakalim, 2-Amino-3-methyl-1-butanol, and 2,4-Dimethyl-1,3-oxazole-5-carboxylic acid showed significant up-regulation, while caffeinol showed down-regulation. Moreover, the differential metabolites were mainly annotated in following pathways; neomycin, kanamycin and gentamicin biosynthesis, two-component system, benzoate degradation, choline metabolism in cancer, terpenoid backbone biosynthesis, plant hormone signal transduction, tryptophan metabolism, acarbose and validamycin biosynthesis, limonene and pinene degradation. Zhang et al. (2023) also reported the significant alterations in metabolites of *S. frugiperda* in response to *B. bassiana* infection. It was found that these differential metabolites were mainly amino acids and their derivatives, lipids, carbohydrates and organic heterocyclic compounds, related to amino acid, lipid, nucleotide, and carbohydrates metabolism^80^.

Existing literature validated that sugars serve as vital source of carbohydrate based energy and are crucial for the growth and development of insects^1^. From the fig. 8a, it is cleared that pyruvate (FC=0.36), pyruvic acid (FC=0.15), succinic acid (FC=0.18) were significantly down-regulated and fumaric acid (FC=1.73) was up-regulated in response to isolate SMH-1 infection. Zhang et al. (2023) indicated that the concentrations of L-tyrosine, L-dopa, arginine, alpha-ketoglutaric acid, cystathionine, D-sedoheptulose-7-phosphate and citric acid were significantly decreased in *B. bassiana* infected larvae of *S. frugiperda* and may lead to oxidative stress, and further converted into the TCA cycle through purine metabolism, arginine biosynthesis, butanoate metabolism, and phenylalanine metabolism^80^. Similarly, the levels of pyruvic acid (FC=0.24) and succinic acid (FC=0.82) were significantly decreased and level of pyruvate (FC=1.77) was increased after seven days of exposure with oral administration of fungal acetone extracts. Further, down-regulation of pyruvic acid (FC=0.12) and fumaric acid (FC=0.66) was found over seven days of feeding with compound dihydrocoumarin, which revealed the perturb TCA cycle of *S. invicta*. Lee et al. (2024) demonstrated that treatment with ethyl formate and low temperature significantly altered the metabolites abundance in *Drosophila suzukii*, the metabolites were mainly annotated into biosynthesis of amino acids, nucleotides and cofactors. Additionally, it found significant alterations in TCA cycle, and the levels of purine and pyrimidine classes of metabolites^81^. Xia et al. (2023) also noted the significantly altered metabolic response of *S. litura* to coumarin treatment, it was found that 391 differentially expressed metabolites mainly related to purine metabolism, amino acid metabolism^75^. Moreover, from 0-24 hours after treatment, 352 differential metabolites involved with ATP-binding cassette (ABC) transporters and amino acid metabolism, which further altered into TCA. Our findings indicated that isolate SMH-1 (conidial pathogenicity, acetone extracts and dihydrocoumarin efficacy) could induce such impactful alterations in the levels of those metabolites, which expected to affect the energy supply mechanism of *S. invicta*.

Taken together, this study demonstrates the biocontrol potential of *C. echinulata* SMH-1 against *S. invicta*, with bioassays confirming 100% mortality via conidial and acetone extract exposure.

Light/electron microscopy revealed fungal infection dynamics, including cuticular penetration and hyphal proliferation. Six metabolites, notably dihydrocoumarin, exhibited significant toxicity, disrupting mitochondrial bioenergetics and detoxification pathways. Metabolomics identified key biomarkers and perturbed pathways (e.g., citrate cycle), elucidating mechanisms underlying *S. invicta* susceptibility. While results advocate SMH-1-derived compounds as eco-friendly alternatives to synthetic insecticides, further validation of dose-response relationships and non-target safety is essential. This work validates metabolomics-driven discovery of entomopathogenic fungal bioactives, advancing sustainable IPM strategies against invasive species. Future research should prioritize field trials and mechanistic synergies to optimize biocontrol efficacy.

## 4. Material and methods

### 2.1 Isolation, identification and culture of entomopathogenic fungal strain

The fungal isolate SMH-1 was obtained from a collapsed *S. invicta* nest located in the hilly terrain adjacent to Yunnan Agricultural University, Kunming, China (25.06025°N, 102.70040°E; elevation: 1,912.90 m). Primary isolation was performed on potato dextrose agar (PDA; Himedia, India) following Qiu et al. (2014), with modifications^36^: PDA medium contained 20 g sucrose, 200 g autoclaved *Solanum tuberosum* extract, and 20 g bacteriological agar (Oxoid, UK) per liter. Sub-culturing was conducted across five generations to ensure genetic stability. Plates were incubated at 25 ± 2°C, 65 ± 5% relative humidity (RH), and a 12:12 h light:dark photoperiod (LED intensity: 2,000 lux) for 10 days. Morphological identification was performed by Dr. Deqiang Qin using lactophenol cotton blue staining, confirming *C. echinulata* via unispored sporangia and pedicellate vesicles^21^. Molecular identification utilized ITS primers (ITS1: 5 ′-TCCGTAGGTGAACCTGCGG-3 ′; ITS4: 5 ′-TCCTCCGCTTATTGATATGC-3 ′) with genomic DNA extracted via HiPurA Fungal DNA Kit (Himedia, Germany) and PCR amplification as per Hussain et al. (2021)^37^.

### 2.2 S. invicta collection and rearing

Colonies of *S. invicta* were excavated using the Banks et al. protocol^38^, transferred to polyethylene containers (32 × 30 × 30 cm) lined with talcum powder barriers, and maintained in acrylic rearing chambers (40 × 25 × 18 cm) under controlled laboratory conditions (25 ± 2°C, 65 ± 5% RH, 12:12 h photoperiod). Colonies were provisioned with 20% (w/v) sterile sucrose solution, protein-rich minced pork sausage (Jinluo, China), and live *Tenebrio molitor* larvae (third instar) on a 48–72 h rotational schedule. Fluon® PTFE (AGC Chemicals, Japan) barriers prevented escape, and environmental parameters were regulated using Saifu PRX-450A climate chambers.

### 2.3 Conidial suspensions preparation for pathogenicity tests

Conidial suspensions were prepared in sterile 0.05% Tween-80 (v/v; Sigma-Aldrich, USA) via logarithmic dilution (1×10⁴–1×10⁸ conidia/mL). For LC_50_ (50% lethal concentration) assays, five concentrations (1×10⁴–1×10⁸ conidia/mL) were tested, while temperature-and size-dependent virulence assays utilized 5.35×10⁷ and 1.03×10⁷ conidia/mL for major (>4 mm) and minor (<4 mm) workers, respectively. Conidia were harvested using camel-hair brushes, homogenized via vortex agitation (VELP Scientifica, Italy), and quantified using a Qiujing® Improved Neubauer hemocytometer (Shanghai Jingjing Biochemical, China) under a BA310-T stereomicroscope (Motic®, China) at 400× magnification.

### 2.4 Ant bioassays

Pathogenicity assays followed Wei et al. (2021) with modifications^39^: cohorts of mixed-sized workers (n = 20 per replicate) were immersed in conidial suspensions (2–3 s), blotted on sterile filter paper (Whatman Grade 1), and transferred to acrylic arenas (9 × 8 × 5 cm) provisioned with 1.5 mL Eppendorf tubes containing 20% sucrose solution. Control groups received 0.05% Tween-80 only. Mortality was recorded daily for 14 days, with cadavers transferred to humidity chambers (70% RH, 25°C) to confirm mycosis via sporulation. LC_50_ values were calculated using probit analysis (SPSS v26, IBM), and worker polymorphism was classified via digital caliper measurements (Mitutoyo, Japan) as per Park et al. (2022) protocol^40^.

### 2.5 Light and Electron microscopic observations

Infection dynamics were analyzed using a KEYENCE VHX-5000 digital microscope (200×; KEYENCE America, USA) and FEI Quanta 450 FEG SEM (Thermo Fisher Scientific, USA). For SEM (scanned electron micrographs), cadavers were fixed in 4% glutaraldehyde (24 h, 4°C), dehydrated in ethanol (30–100%), critical-point dried (Leica EM CPD300), and sputter-coated with Au-Pd (10 nm). Samples were imaged at 1,000–5,000× magnification under 15 kV acceleration voltage, following Hussain et al. (2021)^37^.

### 2.6 Extraction of the fungal material and its efficacy bioassay

Acetate extracts were prepared by submerging 22-day-old SMH-1 cultures (25 ± 2°C, 65 ± 5% RH) in HPLC-grade acetone (1:1 w/v) for 7 days with orbital shaking (120 rpm). Crude extracts were filtered (Whatman No. 1), concentrated via rotary evaporation (40°C, 200 mbar; Büchi R-300), and serially diluted (62.5–1,000 mg/L) in 10% DMSO-methanol. Bioassays utilized 1:9 (v/v) extract-sucrose mixtures in Eppendorf tubes, replaced every 48 hours^41^. Mortality was recorded under standardized conditions (25 ± 2°C, 65 ± 5% RH).

### 2.7 Toxicity test of selected compounds

Six compounds (coumarin, dihydrocoumarin, scopoletin, scopolin, 7-hydroxycoumarin, 5-methoxysporalen; Shanghai Macklin Biochemical, China) were tested at 1–100 mg/L (or μL/L for dihydrocoumarin). Stock solutions (20 mL) were prepared in acetone, diluted in sucrose solution (1:9 v/v), and administered via “water tube” method^42^. Controls received acetone-sucrose (1:9 v/v). Mortality was assessed under identical environmental conditions.

### 2.8 UPLC-MS based metabolomics profiling of isolate SMH-1

#### 2.8.1 Instruments, Reagents and Materials

Metabolite profiling utilized a Waters UPLC Acquity I-Class PLUS/Xevo G2-XS QTOF system (Waters, USA) with an HSS T3 column (1.8 µm, 2.1 × 100 mm). Solvents (methanol, acetonitrile, formic acid) were LC-MS grade (Merck, Germany).

#### 2.8.2 Sample preparation

Mycelia (50 mg) were homogenized in methanol: acetonitrile: H_2_O (2:2:1 v/v) with 20 mg/L L-2-chlorophenylalanine (internal standard), vortexed (30 s), ground (45 Hz, 10 min), and sonicated (10 min, ice bath). After centrifugation (12,000 × g, 15 min, 4°C), supernatants were vacuum-dried, reconstituted in acetonitrile: H_2_O (1:1 v/v), and filtered (0.22 µm) prior to analysis^43,44^.

#### 2.8.3 UPLC-MS conditions

Chromatography employed 0.1% formic acid (A) and 0.1% acetonitrile-formate (B) with gradient elution: 0–0.25 min (2% B), 0.25–10 min (2–98% B), 10–13 min (98% B), 13.1–15 min (2% B).

MS parameters: ESI ± 2,500 V, cone voltage 30 V, source temperature 100°C, desolvation temperature 500°C, m/z 50–1,200.

### 2.9 LC-QTOF based metabolomics profiling of Solenopsis invicta against different treatments

#### 2.9.1 Sample preparation and metabolites extraction

Treated ants were flash-frozen (-80°C), lyophilized, and homogenized in methanol: acetonitrile: H_2_O (2:2:1 v/v). Post-centrifugation, supernatants were processed as above^43,44^.

#### 2.9.2 UPLC-MS (LC-QTOF) conditions

Chromatographic and MS parameters mirrored fungal profiling. Data were acquired in MS-e mode (2–40 V collision energy) using MassLynx v4.2.

#### 2.9.3 UPLC-MS data processing (For both Tests)

Raw data were processed via Progenesis QI (Waters) with METLIN and HMDB alignment. Multivariate analysis (PCA, OPLS-DA) and pathway enrichment (KEGG, lipidmaps) were performed using ropls (R v4.2.1). Differential metabolites were filtered (VIP >1, p <0.05, FC >1)^45^.

### 2.10 Statistical analysis

Mortality data were analyzed via one-way ANOVA (SPSS v26) with Tukey’s HSD post hoc tests (p <0.05). Survival curves were plotted using GraphPad Prism v8.0.

## Acknowledgments

Not applicable.

## Ethics approval

No ethical approval is required.

## Author contributions

MH performed experiments, data analysis and wrote manuscript. XQ helped in review and methodology. XG technical help and resources provision. GW supervised the project and provide funding. DQ funding, supervised, and help in drawing figures. All authors read and approved the manuscript.

## Availability of data and material

The original contributions presented in the study are included in the article/Supplementary Materials. The ITS sequences generated in this study can be found at NCBI with accession number (SUB14919727). Further inquiries can be directed to the corresponding authors.

